# Assemblytics: a web analytics tool for the detection of assembly-based variants

**DOI:** 10.1101/044925

**Authors:** Maria Nattestad, Michael C Schatz

## Abstract

**Summary:** Assemblytics is a web app for detecting and analyzing structural variants from a *de novo* genome assembly aligned to a reference genome. It incorporates a unique anchor filtering approach to increase robustness to repetitive elements, and identifies six classes of variants based on their distinct alignment signatures. Assemblytics can be applied both to comparing aberrant genomes, such as human cancers, to a reference, or to identify differences between related species. Multiple interactive visualizations enable in-depth explorations of the genomic distributions of variants.

**Availability and Implementation:** http://qb.cshl.edu/assemblytics, https://github.com/marianattestad/assemblytics

**Supplementary information:** Supplementary data are available at *Bioinformatics* online.

## 1 Introduction

*De novo* genome assembly is becoming increasingly tractable on large genomes due to advances in long-read sequencing and mapping. This is leading to a greater quality and quantity of reference genomes across the tree of life (Lee, et al., 2014; Roberts, et al., 2013). Researchers can now sequence and assemble the genomes of several related strains or species in order to compare them. This is a vast improvement over more common resequencing approaches where sequencing reads are aligned to a single reference genome, often allowing only SNPs or short indels to be identified. Now that increasing numbers of high-quality genome assemblies are available, there is a need to detect the large structural variants that mark important differences between these genomes. For example, there may be more than 10,000 structural variations representing mega-bases of genetic diversity present per human genome (see below). Assemblytics builds on the innovations of the whole genome alignment suite MUMmer (Kurtz, et al., 2004) in order to detect and analyze these variants.

## 2 Methods

Assemblytics analyzes the alignments from MUMmer’s *nucmer* program to identify high-confidence structural variants in each sequence (contig) in the sample relative to a reference or another *de novo* assembly. It begins by loading the *nucmer* alignments into an interval tree to quickly identify all overlapping alignments with respect to the sample. It then filters the alignments to report those with at least a minimum amount of unique contig sequence anchor (default: 10kbp) contained in no other alignments of that contig. This is similar to the filtering performed by *delta-filter* component of *dnadiff* (Phillippy, et al., 2008), although guarantees uniqueness of the alignments while *dnadiff* may select equally matching repetitive alignments arbitrarily (**Supplementary Note 1**).

The variant identification algorithm then considers each pair of consecutive alignments along a sample contig, determining variant presence and class by the spacing and orientation between these alignments. This identifies all variants at least 50 bp long (the standard definition of a structural variation) up to a maximum of 10 kbp in size, with this maximum adjusted to match the size of the unique sequence anchor. This prevents translocations and complex variants from being interpreted as indels. Figure 1A illustrates the differences between variant classes. For insertions and expansions the contig contains more sequence than the reference, whereas for deletions and contractions the contig contains less sequence than the reference. Insertions and deletions are characterized by a defined breakpoint (less than 50bp overlap or gap) on one side. Tandem variants are characterized by overlapping alignments (over 50 bp) on either side or both. Repeat variants are characterized by gap in alignment (over 50 bp) on both sides. The individual alignments are also scanned to detect insertions or deletions of at least 50 bp that were fully spanned by the alignment. Finally variant classes, size distributions, and genomic coordinates of all variants are summarized through plots and tables (Figure 1B & C). **Supplementary Note 3** provides more details on the web interface.

**Figure 1.**
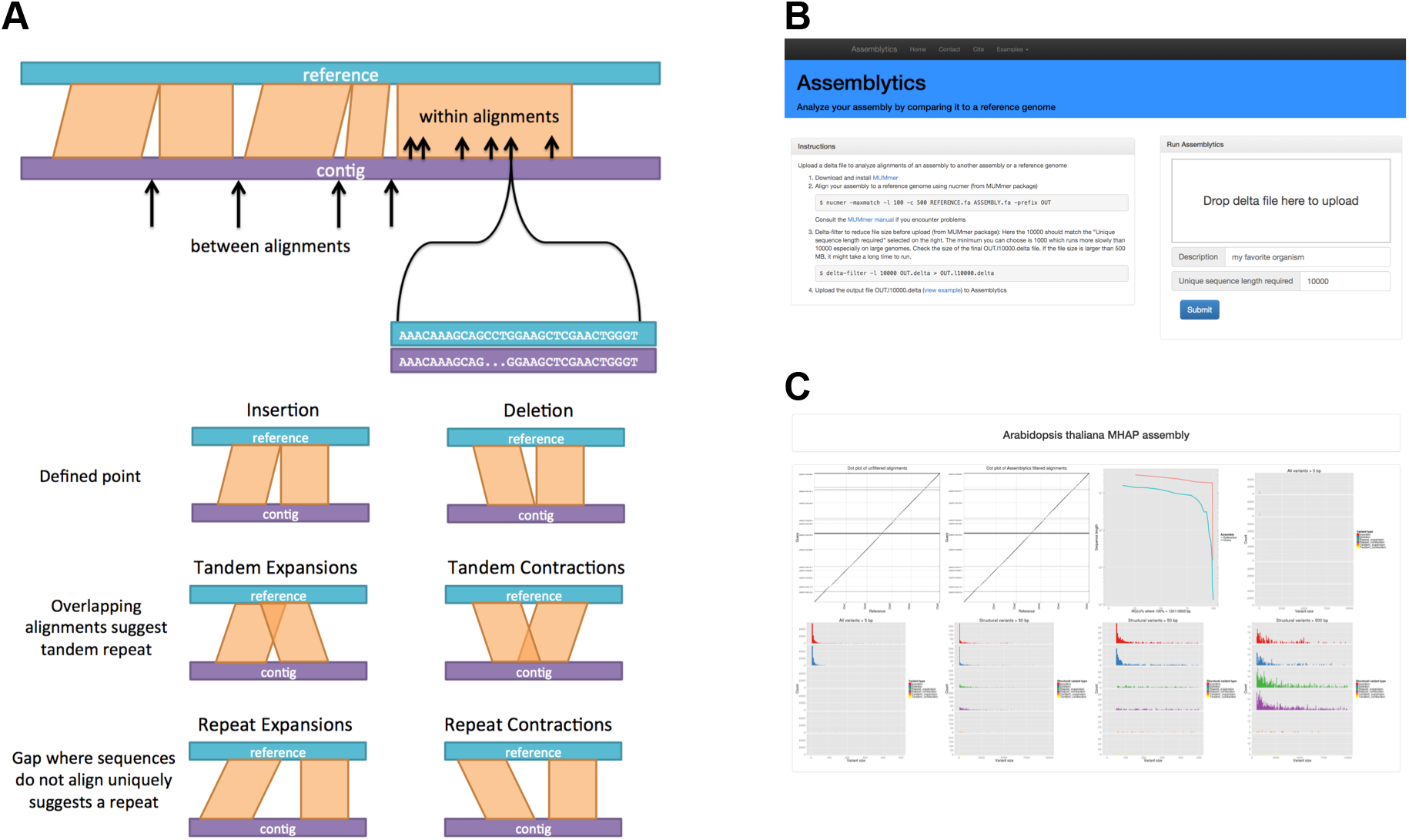
**(A)** Schematic illustrating how variants are called between consecutive alignments of a contig to the reference as well as within alignments. Each variant class is characterized by the degree of gap and/or overlap between alignments. **(B,C)** Screen shots of the Assemblytics web interface including output plots. Additional screen shots in **Supplementary Note 3** show summary tables, variant file preview, and ability to download all data including the Assemblytics unique anchor filtered delta file.

## 3 Results

We first evaluated the accuracy of the variation detection by applying Assemblytics to analyze simulated structural variations within the human genome **(Supplemental Note 2)**. To do so, we created a modified version of the human genome with simulated insertions and deletions embedded at known positions, and then aligned both the original unmodified reference genome and the human genome assembled by MHAP presented by Berlin, et al. (2015) to this modified reference. Our results show that Assemblytics is able to correctly identify the vast majority of variants present (90.9% to 99.9% recall) and with very low false positive rates (0.29% to 0.40%).

We next applied Assemblytics to all five de novo assemblies (human, *D. melanogaster, A. thaliana, S. cerevisiae*, and *E. coli)* presented by Berlin et al (2015) to their respective reference genomes **(Supplementary Note 4)**. In each case, the genome was analyzed in less than 5 minutes and the full interactive results are available on the Assemblytics website. All assemblies showed a large number of structural variations, proportional to the size of the genome, and the sizes of the events approximated a log-normal distribution with a long tail of large events. Drosophila was a notable exception with a relatively small number of variants identified, with only 204 compared to 2,501 for A. *thaliana* of similar size. This was because the assembly was derived from the same inbred population as the original reference (ISO1). In the case of the human analysis, Assemblytics reported 11,206 structural variations spanning over 7.0Mb. This included a noticeable enrichment for ∼320 bp insertions and deletions which we characterized as novel Alu variants.

## 4 Discussion

Assemblytics allows researchers to take advantage of high-quality genome assemblies for detecting structural variation between species or even between aberrant and normal genomes in human disease. Here we applied Assemblytics to study 5 *de novo* assemblies produced using MHAP from PacBio reads, although it is capable of exploring assemblies and structural variants produced by any algorithm and sequencing technology. The accuracy of variant calls depends on the qualities of both assembly and reference genome as well as the accuracy of the whole genome alignments. A key advantage of Assemblytics is unique length filtering which disregards alignments that are not anchored in a significant amount of unique sequence. This provides a conservative filter, similar to requiring high mapping quality for read alignment, something that was not previously available for genome-genome alignments.

## Acknowledgements

We thank Jason Chin, Adam Phillippy, Art Delcher, W. Richard McCombie, Fritz Sedlazeck, and James Gurtowski for their helpful discussions and testing.

## References

Berlin K., et al. Assembling large genomes with single-molecule sequencing and locality-sensitive hashing. Nat Biotechnol 2015; 33(6): 623–630.

Kurtz S., et al. Versatile and open software for comparing large genomes. Genome Biol 2004; 5(2): R12.

Lee H., et al. Error correction and assembly complexity of single molecule sequencing reads. bioRxiv 2014.

Phillippy A.M., Schatz M.C. and Pop M. Genome assembly forensics: finding the elusive mis-assembly. Genome Biol. 2008; 9(3): R55.

Roberts R.J., Carneiro M.O. and Schatz, M.C. The advantages of SMRT sequencing. Genome Biol. 2013; 14(7): 405.

